# Identification of Genomic Insertion and Flanking Sequences of the Transgenic Drought-tolerant Maize Line “SbSNAC1-382” using the Single Molecular Real-Time (SMRT) Sequencing Method

**DOI:** 10.1101/858951

**Authors:** Tingru Zeng, Dengfeng Zhang, Yongxiang Li, Chunhui Li, Xuyang Liu, Yunsu Shi, Yanchun Song, Yu Li, Tianyu Wang

**Affiliations:** Institute of Crop Sciences, Chinese Academy of Agricultural Sciences, Beijing, China

**Keywords:** transgenic maize, flanking sequence, fosmid library, SMRT sequencing

## Abstract

Safety assessment of genetically modified (GM) crops is crucial in the phase of product development before the GM crops are put on the market. Characteristics of flanking sequences of exogenous insertion sequences are essential for the safety assessment and marking of transgenic crops. In this study, we used the methods of genome walking and whole genome sequencing (WGS) to identify the flanking sequence characteristics of a *SbSNAC1* transgenic drought-tolerant maize line “SbSNAC1-382”, but both of the methods failed. Then, we constructed a genomic fosmid library of the transgenic maize line, which contained 4.18×10^5^ clones with an average insertion fragment of 35 kb, covering 5.85 times of the maize genome. Subsequently, three positive clones were screened by pairs of specific primers and one of the three positive clones was sequenced by using the Single Molecule Real-Time (SMRT) sequencing technology. More than 1.95 Gb sequence data (∼10^5^ × coverage) for the sequenced clone was generated. The junction reads mapped to the boundaries of T-DNA and the flanking sequences in the transgenic line were identified by comparing all sequencing reads with the maize reference genome and the sequence of transgenic vector. Furthermore, the putative insertion loci and flanking sequences were confirmed by PCR amplification and Sanger sequencing. The results indicated that two copies of the exogenous T-DNA fragments were inserted in the same genomic site. And the exogenous T-DNA fragments were integrated at the position of Chromosome 5: 177155650 to 177155696 in the transgenic line 382. Herein, we have demonstrated the successful application of the SMRT technology for the characterization of genomic insertion and flanking sequences.

## Introduction

Since genetically modified (GM) crops were first introduced in the U. S. in the mid-1990s, they have become widely adopted by growers of many countries in the world [1]. In 2017 alone, 189.8 million hectares of GM crops were planted worldwide [2]. It is an international consensus that GM crops could be commercialized after they are proven to be safe. As a result, extensive testing and comprehensive analyses of transgenic lines with excellent objective traits are necessary for biosafety assessment before being approved and entering into market. Among these, molecular characterization of GM crops at the chromosomal level including insertion sequences, sites, copy numbers and flanking sequences is essential for the safety assessment and specific detection of GM crops [3, 4]. Furthermore, identification of T-DNA flanking sequences of GM crops and the development of specific detection methods are useful for breeding program, and important for bio-risk management to ensure food, feed and environmental safety [5, 6].

Traditionally, exogenous fragments flanking sequences of transgenic plants are obtained by various PCR-based methods according to the T-DNA sequence information [7-10]. Among them, thermal asymmetric interlaced PCR (TAIL-PCR) and genome walking are often used to isolate and clone T-DNA flanking sequences [9, 10]. Using the TAIL-PCR method by sequencing, several T-DNA flanking sequences were identified and characterized in transgenic maize [11], soybean [12], cotton [13], and alfalfa [14]. However, these PCR-based methods are laborious and expensive.

With the emergence and rapid development of next-generation sequencing (NGS) technology over the past few years, molecular characterization of insert locations, copy numbers, integrity, and stability of transgenic crops can be implemented in a relatively short time and at acceptable cost [15]. Up to now, a number of the flanking sequences of exogenous genes in GM plants such as *Arabidopsis* [16], rice [17], soybean [6], and maize [18] have been identified by the NGS method.

Taken together, both the PCR-based method and the NGS technology enabled us to successfully characterize both single and stacked transgenic events [15]. However, these approaches are difficult to identify all insertion loci and their flanking sequences of transgenic events with complex genome sequences or intricate modifications or rearrangements of exogenous fragments [6, 19].

Maize is one of the most important crops in the world and 31% of the GM crops’ growing area planted annually in the world are GM maize [1]. It is important to evaluate the safety of the GM maize, especially to identify the flanking sequence of exogenous genes in the GM maize. However, maize has larger genome and more repetitive sequences compared with soybean, cotton and rice, and it is difficult to identify flanking sequences of inserted genes [20]. In addition, transgenic maize events may often contain a part of or the entire vector backbone. In other cases, a partial copy of T-DNA inserts and the connection takes place outside the expected boundary [21, 22]. Therefore, for the acquisition of flanking sequences integrated into larger genomes and complex insertion fragments, the accurate flanking sequences can often be found by constructing DNA libraries. Turning genomes into countless fragments by physical or biological means cloned in fosmid or BAC vectors were a mainstay of genome projects during the Sanger-based sequencing era [23, 24]. Compared with other libraries, fosmid libraries have the advantages of short cloned fragments (about 40 kb), single copy insertion and easier to generate [25]. Because inserts in the fosmid libraries are generated randomly by ultra-sound rather than by enzymatic digestion, inserts in the fosmid libraries can avoid potential clone biases. It is suitable for physical mapping, gene cloning and chromosome mapping of gene fragments [6, 26]. Recently emerged single-molecule based NGS technology generate longer reads (20 kb) at increased coverage depth and is particularly important in resolving the challenges in characterization of transgenic events with insert locations in repetitive and low complexity regions of a genome [27]. As a result, using the fosmid libraries and the single-molecule based NGS technology might be suitable for identifying T-DNA flanking sequences of transgenic lines with complex genome sequences or intricate modifications or rearrangements of exogenous fragment.

Recently, we developed one transgenic line “SbSNAC1-382” by over-expression of *SbSNAC1* from sorghum, which conferred drought tolerance without a cost of crop productivity under well-watered conditions. Southern blots confirmed that the transgenic line SbSNAC1-382 was a two-copy insertion event, and the two copies might be inserted at the same genome location. In order to obtain the flanking sequence of the target gene of the transgenic maize event, after the failure of the genome walking method and the whole genome sequencing method, the single molecule real-time sequencing was used to identify the accurate flanking sequences of the inserted gene. Molecular characterization of the drought-tolerant transgenic maize at nucleic acid level will provide precise information for regulatory submissions and facilitate utilization of the line in future breeding program.

## Materials and methods

### Plant materials

*SbSNAC1* with *Bst*E II and *Nco***I** enzymatic restriction sites were recombined into the pCAMBIA3301 vector under control of the cauliflower mosaic virus (CaMV) 35 S promoter, resulting in 35S::*SbSNAC1* constructs (S1 Fig). The constructed vector was transferred into maize hybrid HiII by *Agrobacterium*-mediated method. Positive transgenic events backcrossed with the inbred line “Zheng58” for six generations, and the resulting “SbSNAC1-382” was used in subsequent flanking sequence identification.

### DNA isolation and Southern blot analysis

Genomic DNA for leaf samples of the transgenic event and the non-transgenic control was isolated using the CTAB method [28].

Thirty micrograms of the genomic DNA from the transgenic event and the non-transgenic control were digested with the restriction enzymes of *BstE*II and *Nco*I overnight. The resolved genomic DNA was then transferred to the positively charged nylon membranes (Hybond-N^+^, Amersham Pharmacia Biotech) using a model 785 vacuum blotter system (Bio-Rad). The *Bar* amplified fragment (Table 1) labeled by DIG high primer DNA labeling (Roche, Cat. No. 11585614910) and purified using a high pure PCR product purification kit (Roche, No. 11732668001). The DNA blots were prehybridized at 42°C for 1 h in DIG easy hyb granule and then hybridized to denatured DIG-labeled probes for 20 h. The blots were then washed twice with 2×SSC and 0. 1% (w/v) SDS for 15 min each and washed twice with 1×SSC and 0. 1% (w/v) SDS for 15 min each. Immunological detection of the probes was carried out in accordance with the manufacturer’s instructions for the DIG high primer DNA labeling and detection starter kit II.

**Table 1.**
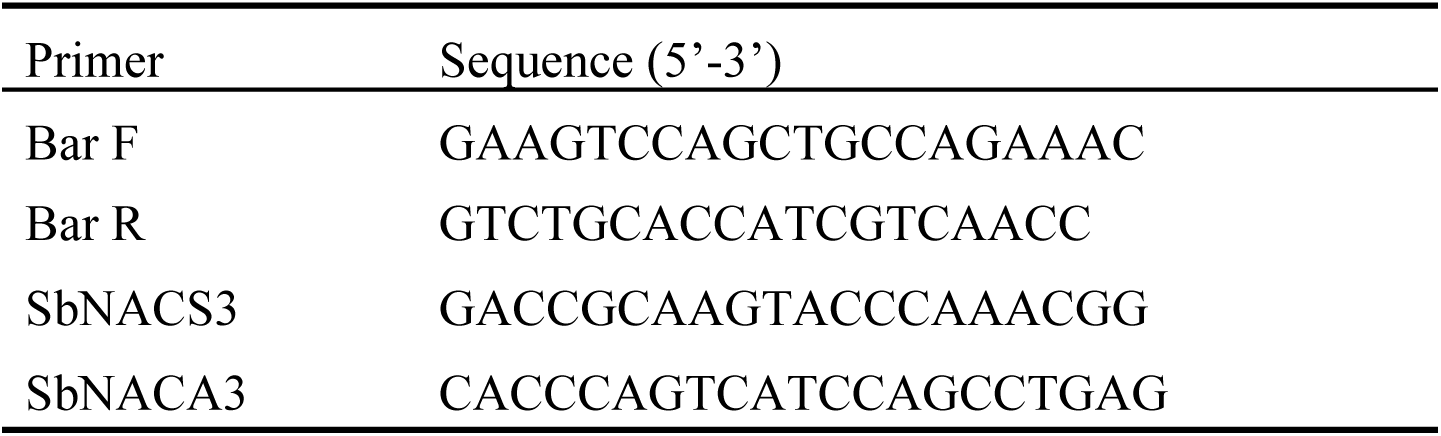
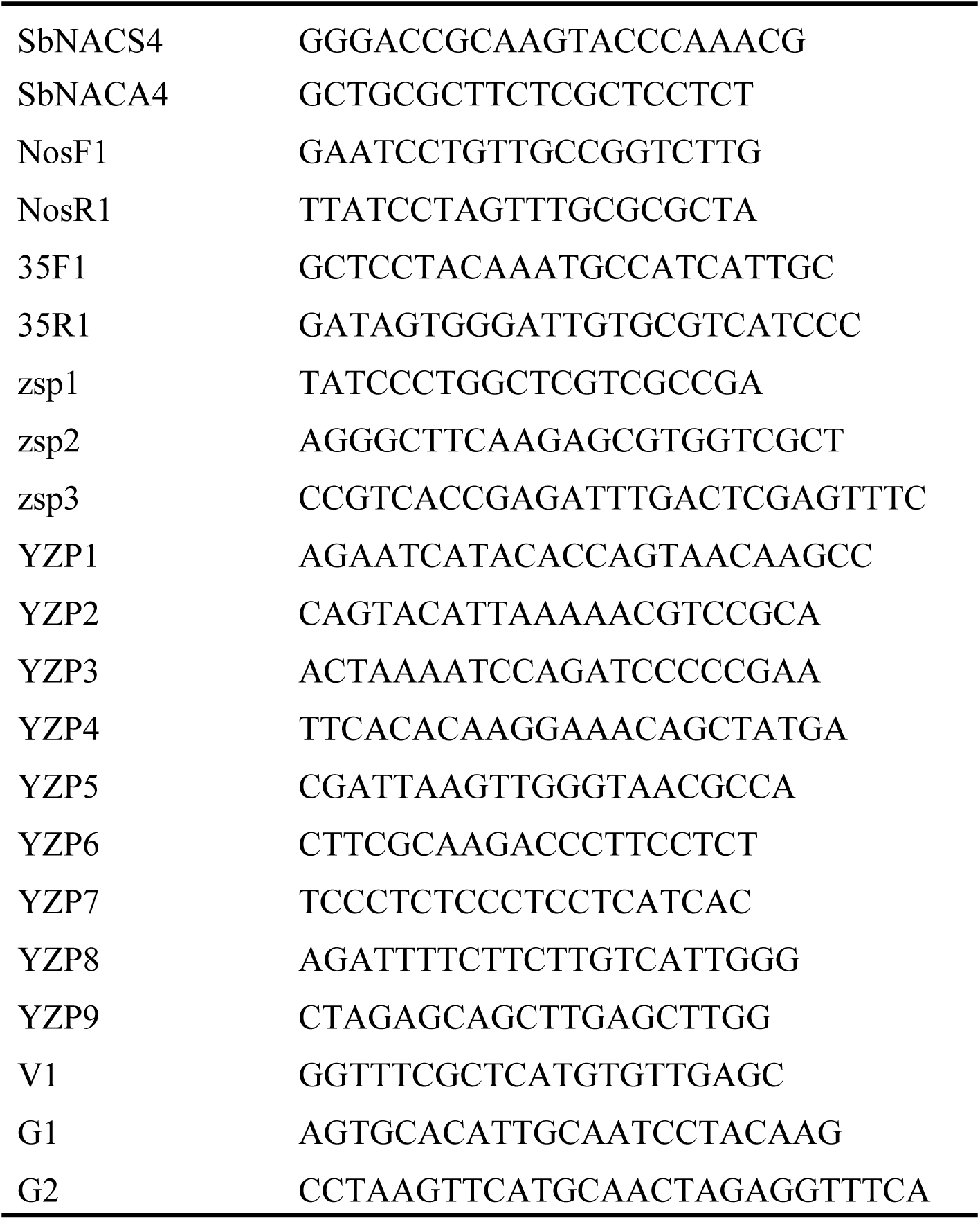
Primers used in this study.

### Genome walking method

The 5’ flanking sequence of the insertion sequence was obtained by the Genome Walking Kit according to the manufacturer instructions (TaKaRa, Dalian, China). The random primers were provided by the Genome Walking Kit and the specific primers designed based on theoretical insertion sequences (first round zsp1; second round zsp2; third round zsp3, Table 1). The specific PCR products were gel purified by using the DNA Gel Extraction Kit (Axygen, USA) and cloned to the pMD-18 vector system (Takara), and then sequenced by the Shanghai Sangon Company.

### Whole genome sequencing

A total of 1.5 μg DNA per sample was used as input material for the DNA sample preparations. Sequencing libraries were generated by using the Truseq Nano DNA HT Sample preparation Kit (Illumina USA) following manufacturer’s recommendations and index codes were added to attribute sequences to each sample. Briefly, the DNA sample was fragmented by sonication to a size of 350 bp, then DNA fragments were end polished, A-tailed, and ligated with the full-length adapter for Illumina sequencing with further PCR amplification. At last, PCR products were purified (AMPure XP system) and libraries were analyzed for size distribution by Agilent2100 Bioanalyzer and quantified using real-time PCR. These libraries constructed above were sequenced by Illumina HiSeq4000 platform and 150 bp paired-end reads were generated with insert size around 350 bp.

### Construction and screening of the fosmid library

DNA was interrupted by ultra-sound and separated by the method of Pulsed Field Gel Electrophoresis (PFGE). DNA fragments from 38-48 kb were recovered and end-repaired the sheared DNA to blunt and 5’-phosphorylated ends. The fosmid library was constructed with the Copy Control™ HTP Fosmid Library Production Kit (Epicenter, USA) using the pCC2FOS™ Vector and EPI300-T1^R^ plating cells.

Three pairs of vector-specific primers were designed to screen positive clones from the fosmid library (SbNACS3/SbNACA3; SbNACS4/SbNACA4; Bar F/Bar R, Table 1). In the initial screening of the library, three pairs of primers were used to detect the library, and a positive colony was obtained. Colony PCR reaction contained 10 μl 2×La Taq Mix (Takara), 1 μl colony, 0.5 μl forward and reverse primer, 8 μl ddH_2_O. The procedure of PCR was as follows: 95°C for 5 min; 95°C for 30 sec; 60°C for 30 sec; 72°C for 30 sec; and a final extension at 72°C for 5 min; 32 cycles. When a positive clone was identified, the positive colony diluted 2 × 10^6^ times with LB liquid media was plated on LB solid medium, and monoclones were picked and subjected to colony PCR.

### Single Molecule Real-Time Sequencing

10 μg of the monoclonal plasmid was extracted and purified. The PacBio libraries were constructed using plasmid that was mechanically sheared to a size of ∼22 kb, using Covaris g-TUBE (Covaris, Inc., Woburn, MA) according the manufacturer’s instructions. PacBio SMRTbell libraries were prepared by ligation of hairpin adaptors at both ends of the DNA fragment using the PacBio DNA template preparation kit 2.0 for SMRT sequencing on the PacBio RS II machine (Pacific Biosciences of California, Inc., Menlo Park, CA). Bluepippin preparation system (SAGE science, Beverly, MA) was used to enrich more than 7 kb fragments in the library. Then, the quality of the library was tested by the Agilent Bioanalyzer 2100 kit (Agilent Technology, Inc., Santa Clara, CA). Sequencing was performed on the PacBio RS II instrument as per the manufacturer’s recommendations.

## Results

### Southern blot analysis of the transgenic line

In order to determine the transgene copy number, a Southern blot analysis were performed by using probes designed to hybridize the *Bar* gene in the T-DNA sequences. The results showed the transgenic line had two copies of insertion of the exogenous sequences when *Hind*III and *Eco*RI were used to digest the DNA of the transgenic line (Fig 1). On the other hand, there was only one band when the DNA of the transgenic line was digested with the restriction endonucleases of *Bgl*II and *Dra*I for which there are no restriction sites in the insertion sequences (Fig 1). As a result, it might be two copies of insertion sequences at the same genomic location of the transgenic maize line.

**Figure 1.**
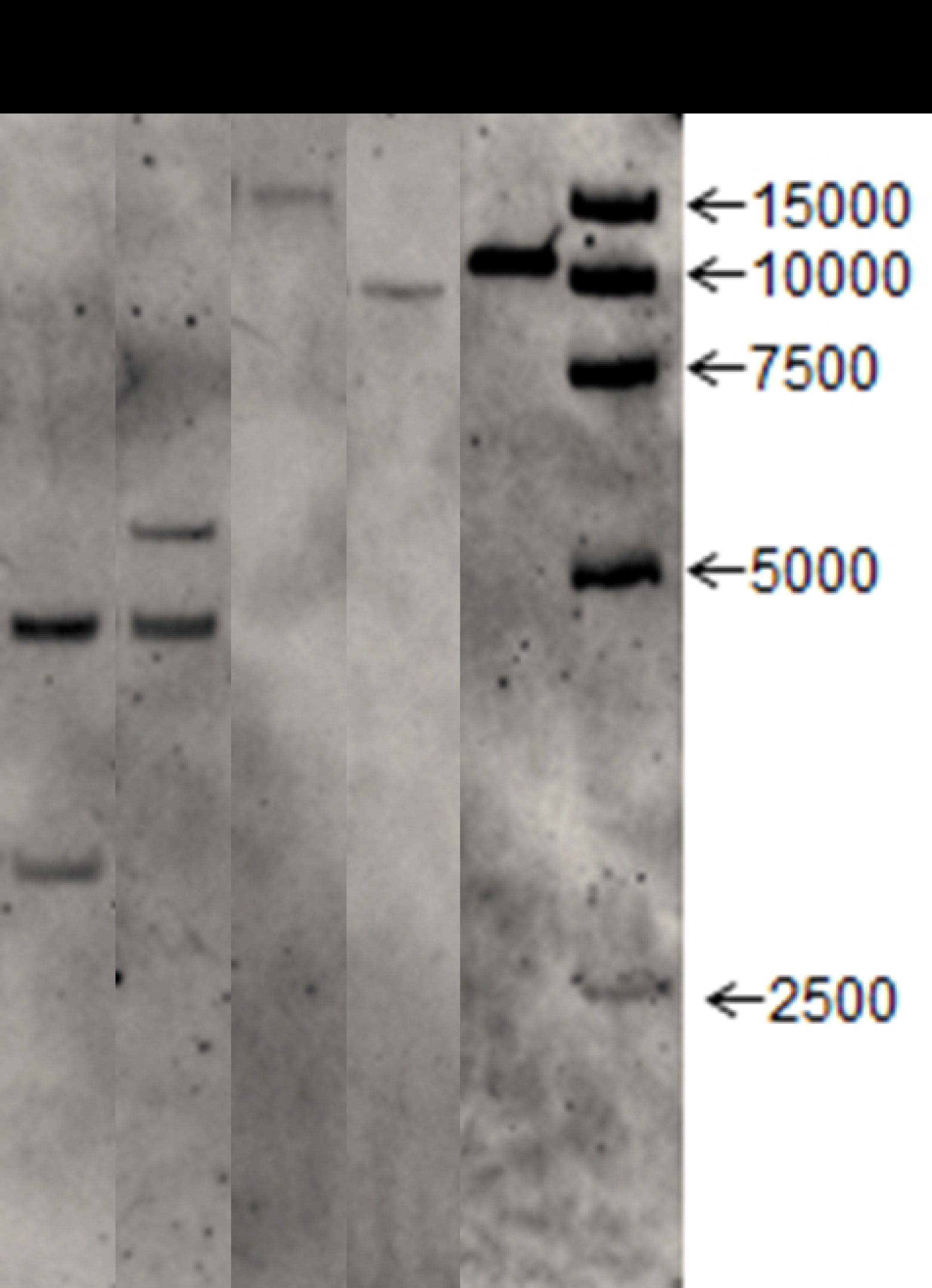
Southern blot analysis of the SbSNAC1-382 line. 1 to 4 digested DNA of the transgenic line by *Hind***III**, *Eco*RI, *Bgl***II** and *Dra*I, respectively; 5, digested plasmids as positive controls, M, marker.

### Genome walking for detecting flanking sequences

Three nested specific primers (zsp1, zsp2, zsp3) were designed according to the sequences adjacent to the T-DNA left border. According to the instructions of the Genome Walking Kit (Takara-Bio, Dalian, China), with nested specific primers and four degenerate primers, three rounds of nested PCR reaction were completed and specific band was obtained (lane 10 of Fig 2). The sequencing results demonstrated that the specific PCR fragment contained 1227 bp in length. By aligning with the maize genome sequence on Maize GDB (www.maizegdb.org) and the T-DNA sequence, it showed that the fragment was made up of 932 bp of non-insert DNA and 295 bp of insert DNA. As expected, the 295 bp inserted DNA was identical to the sequence which was adjacent to the T-DNA left border. The 932 bp of non-insert DNA was identical to the maize genome sequence which is located between 177155650 - 177156582 bp on Chromosome 5. However, the flanking sequence adjacent to the T-DNA right border could not be identified with multiple nested specific primers and degenerate primers using the same method.

**Figure 2.**
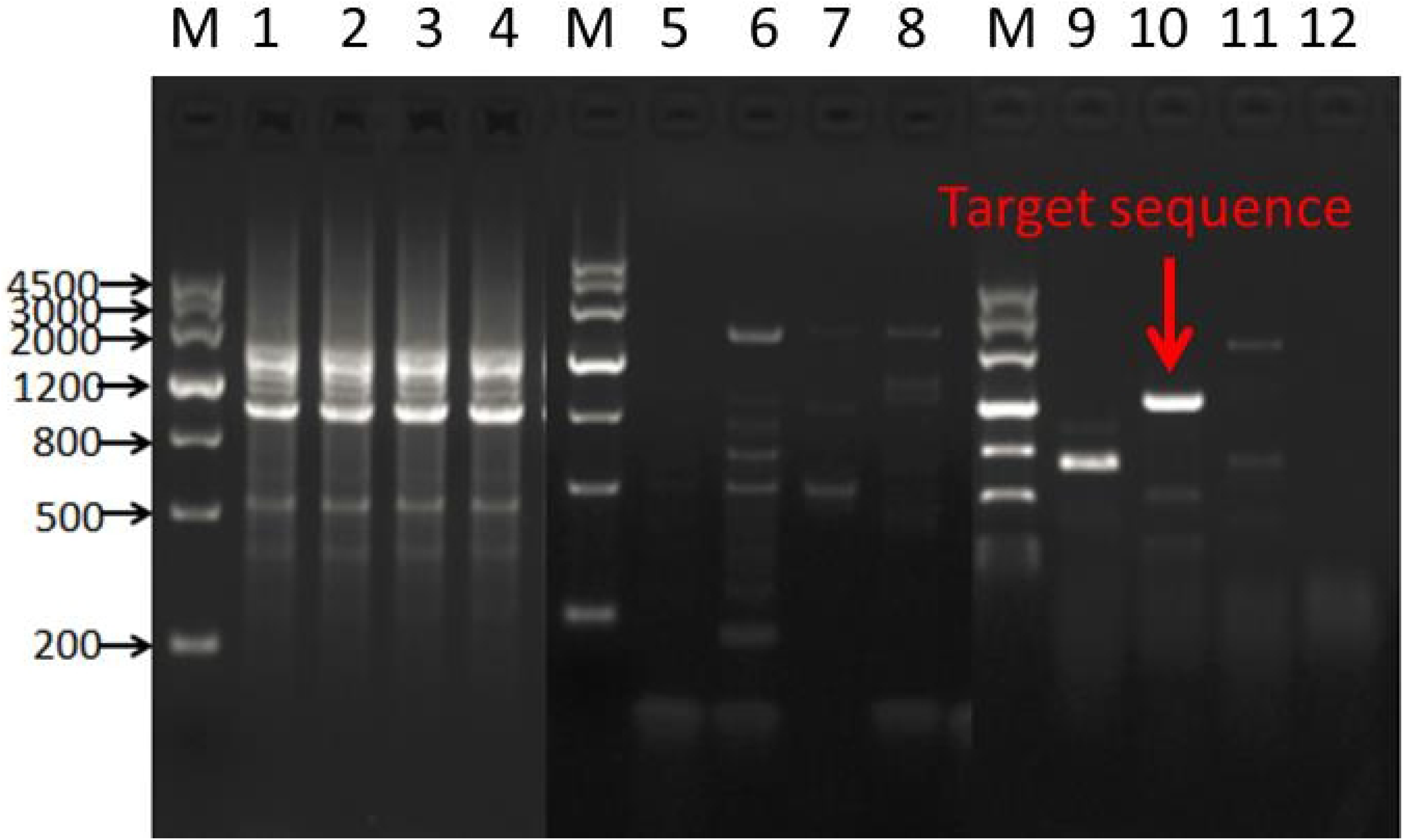
Genome walking results for 5’ flanking sequence. 1-4 lanes are the first amplification results of specific primer zsp1 and degenerate primer AP1-AP4, respectively; 5-8 lanes are the second amplification results of specific primer zsp2 and degenerate primer AP1-AP4, respectively; 9-12 lanes are the third amplification results of specific primer zsp3 and degenerate primer AP1-AP4, respectively; M: marker.

### WGS for detecting flanking sequences

We attempted to use WGS technique to identify flanking sequences on both sides of the insertion sequence. Sequencing libraries were sequenced by Illumina HiSeq4000 platform and 150 bp paired-end reads were generated with insert size around 350 bp. After quality control processing, a total of 144.6 billion clean reads for the transgenic line were obtained (Table 2). Among them, 97. 66% of the reads could be mapped to the reference genome, accounting for ∼64.57 × coverage of the maize genome. Furthermore, about 93.66% of the genome had at least one-fold coverage and 88.57% had at least four-fold coverage. Therefore, the above data indicate that the quality of sequencing was qualified and met the requirements of analysis.

**Table 2.**
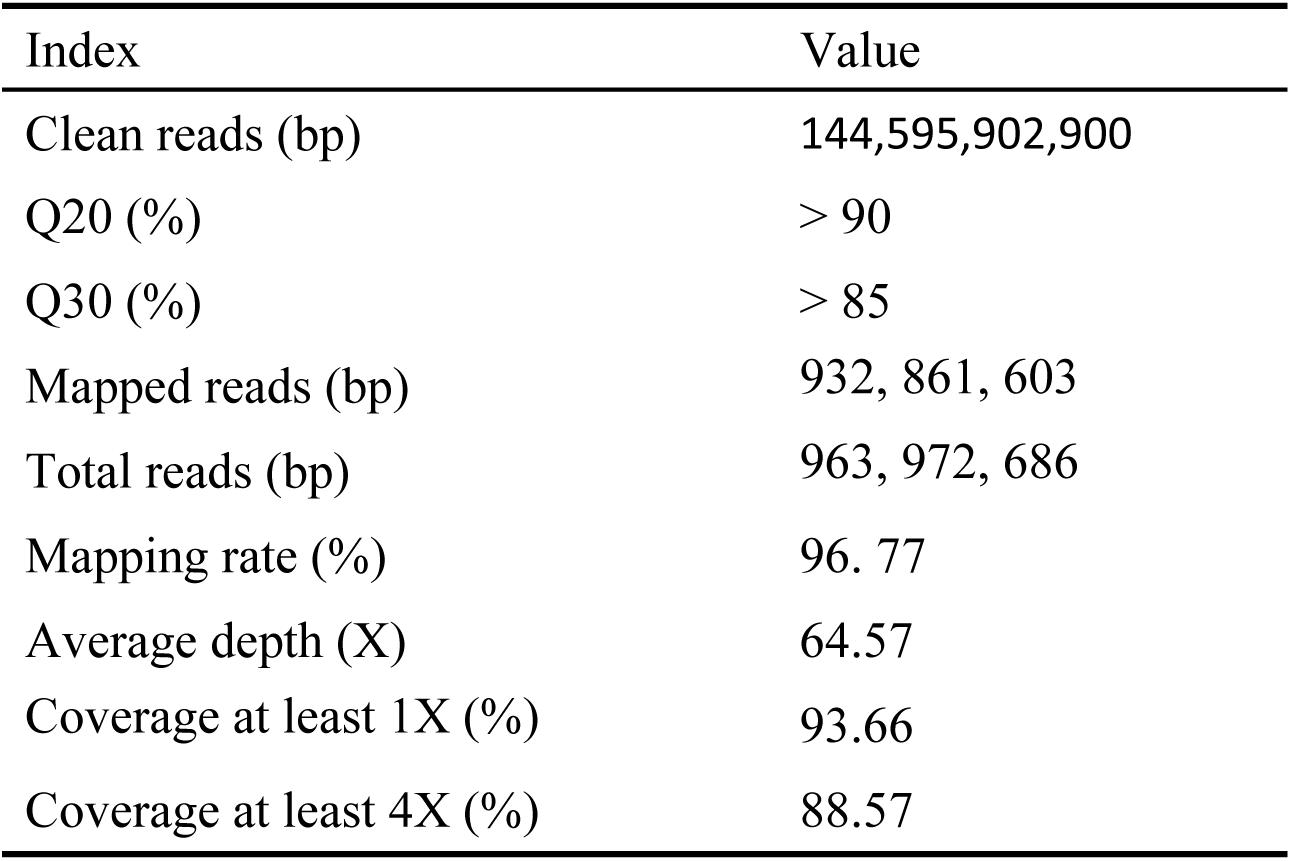
The summary of sequence data of WGS.

| Index | Value |
| --- | --- |
| Clean reads (bp) | 144,595,902,900 |
| Q20 (%) | > 90 |
| Q30 (%) | > 85 |
| Mapped reads (bp) | 932, 861, 603 |
| Total reads (bp) | 963, 972, 686 |
| Mapping rate (%) | 96. 77 |
| Average depth (X) | 64.57 |
| Coverage at least 1X (%) | 93.66 |
| Coverage at least 4X (%) | 88.57 |

In order to identify flanking sequences of putative insertion sites of exogenous fragments, all clean reads were mapped to the sequence of *pCAMBIA3301-SbSNAC1* vector and the maize reference genome. The putative flanking sequence of SbSNAC1-382 line was characterized based on junction reads in which one end of which maps to the vector sequence and the other end to the maize genome. After detailed analysis, five putative flanking sequences were found. One of the five possible flanking sequences was consistent with the Genome Walking’s results. The total length of the fragment was 263 bp. The 150 bp DNA sequence was identical to the sequence adjacent to the T-DNA left border, and the 113 bp DNA sequence was identical to the maize genome. Unfortunately, that the other four putative flanking sequences were not true according to the PCR results. As a result, the flanking sequence adjacent to the T-DNA right border of the SbSNAC1-382 line was still not identified by using the WGS technology.

### Fosmid library construction and positive clone screening

To identify the flanking sequence adjacent to the T-DNA right border of the SbSNAC1-382 line, we constructed a fosmid library of the SbSNAC1-382 line (Takara, Dalian, China), with the recombination rate of 100% (S2 Fig). The original library was diluted and the number of colonies was counted. The library contained about 4.18 × 10^5^ clones, and the average length of the inserted fragments was about 35 kb, which could achieve 5.85 times of the maize genome coverage. According to the Clarke-Carbon formula [29], the probability of screening any gene or sequence from the constructed library was 99. 71%.

In order to screen the target clone from the fosmid library, five pairs of primers were designed (SbNACS3/SbNACA3; SbNACS4/SbNACA4; Bar F/Bar R, NosF1/NosR1, 35F1/35R1, Table 1) according to the T-DNA sequences. According to the results of PCR methods, three positive clones were identified and stored for single-molecule real-time sequencing.

### Single-molecule real-time sequencing and Sanger sequencing

One of the three positive clones screening from fosmid library was selected for sequencing. After the processing of quality control, yielding a total of 1.95 Gb in 100,544 clean reads with a mean length of 9.5 Kb, an N50 length of 12.5 Kb (Table 3).

**Table 3.** Statistics of single-molecule real-time sequencing for plasmid DNA.

| Index | Value |
| --- | --- |
| Polymerase read bases (bp) | 1,949,620,057 |
| Number of polymerase reads | 100,544 |
| Post-filter mean read length (bp) | 19,390 |
| Polymerase read N50 (bp) | 28,065 |
| Polymerase read quality | 0.83 |
| Mean subread length (bp) | 9,509 |
| Subreads N50 (bp) | 12,507 |
| Number of subreads | 204,500 |

To determine the hypothetical insertion sites of exogenous fragments, we constructed a local BLAST data library of single-molecule real-time sequencing data. According to the BLAST results of T-DNA sequences with the local data library, it confirmed the results of Southern blot that the exogenous sequences were composed of two copies of the *SbSNAC1* gene and the *bar* gene at the same maize genomic location. And the flanking sequences of both right border and left border were identified. In order to confirm the flanking sequences of the SbSNAC1-382 transgenic line, specific PCR primers were designed based on the putative genomic sequences and the insertion sequences. When using primer pairs with one primer annealing within putative flanking sequences (YZP2, YZP3, G1, G2, Table 1) and the other annealing to the insertion sequence (YZP1, V1, Table 1), the gel electrophoresis revealed that PCR reactions of primer pairs (YZP1/YZP2; YZP1/YZP3; V1/G1; V1/G2, Table 1) had generated products with single band in the transgenic event 382 while no correct product could be detected from the non-transgenic control of Zheng58 or negative control of water (Fig 3). In addition, YZP3/G2 (Table 1) primers were used to amplify the whole length of the inserted sequence and sanger sequencing of the PCR products showed that the sequence was basically the same as that of the single-molecule sequencing, except for a few bases. Therefore, the exogenous sequence of the SbSNAC1-382 line was integrated at the physical position of Chr. 5: 177,155,650 to 177,155,696 with a 46 bp deletion (Fig 4). Furthermore, the exogenous fragment was inserted into the intergenic region of the maize genome, and no functional genes were interrupted by the inserted sequence.

**Fig 3.**
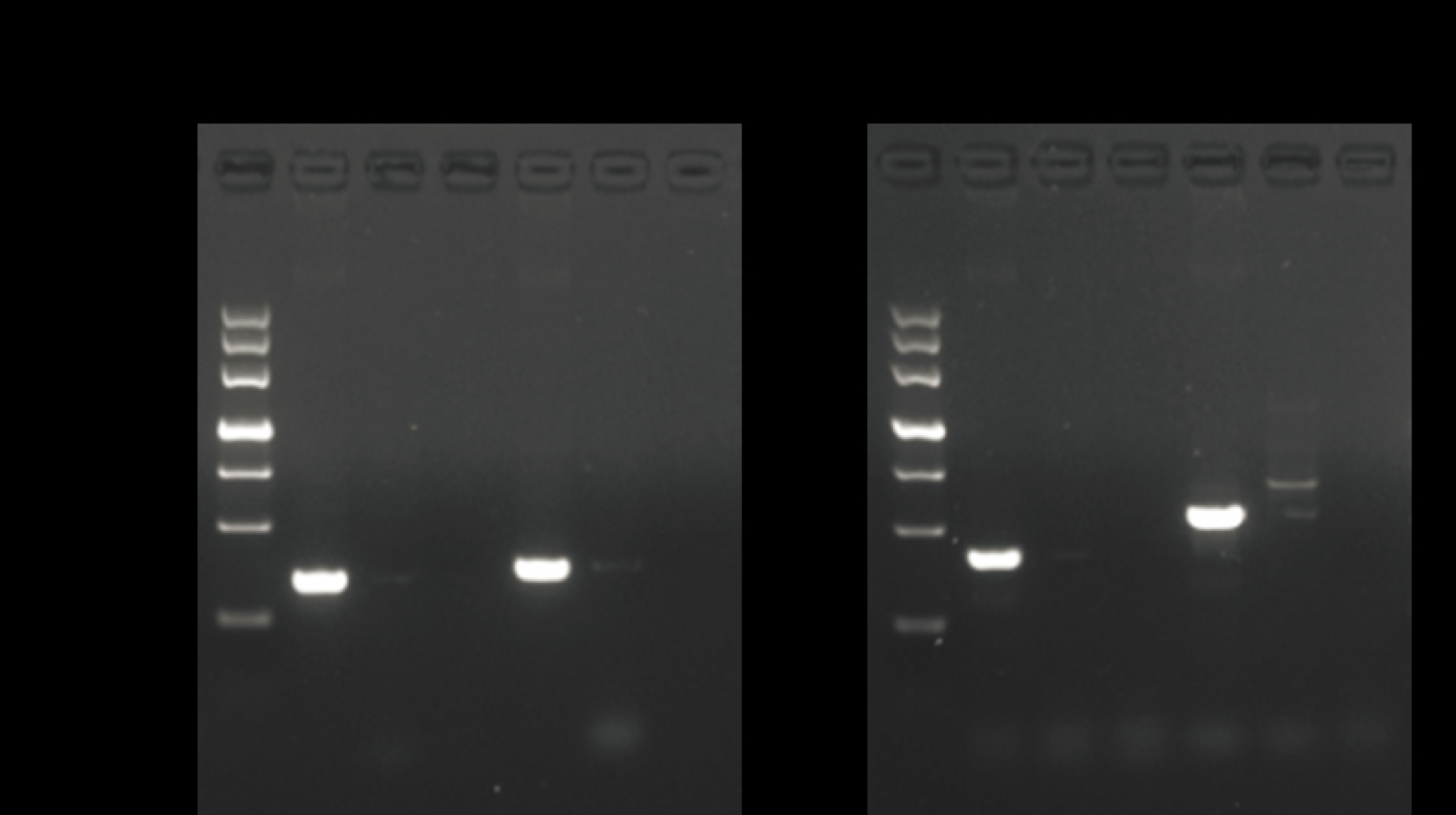
PCR validation of transgenic insertion sites. (A) PCR verification of 5’ end of inserted sequence. 1, 2, 3 and 4, 5, 6 primer YZP1/YZP2 and YZP1/YZP3 amplified in the transgenic line, negative control Zheng58, negative control of water, respectively. M: marker. (B) PCR verification of 3’ end of inserted sequence. 1, 2, 3 and 4, 5, 6 primer V1/G1 and V1/G2 amplified in the transgenic line, negative control Zheng58, negative control of water, respectively.

**Fig 4.**
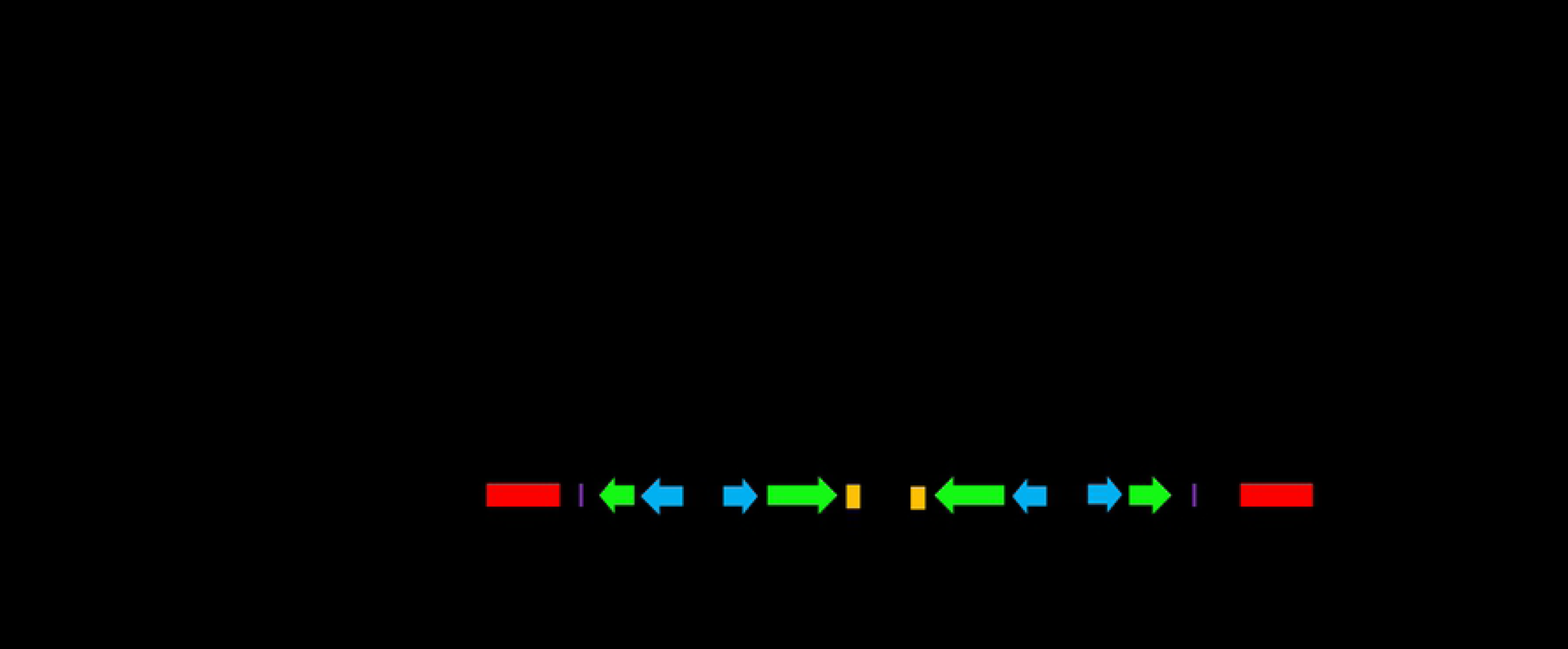
Schematic diagram of insertion loci and flanking sequences of SbSNAC1-382. The numbers under the line of Chr. 5 indicates physical positions on the chromosome. The arrows indicate the position of the validation primer.

In order to verify the results of single-molecule sequencing, we designed a series of primers on the insertion sequence (YZP2/YZP5; YZP4/YZP7; YZP6/YZP9; YZP8/G1, Table 1). The PCR products obtained by these primers were sequenced and compared with the results of single-molecule sequencing. It was found that the two sequencing results were basically the same, showing that single-molecule sequencing had a high accuracy. Further analysis of the structure of the insertion sequence revealed that the exogenous sequence contained two insertion sequences with tandem repetition and opposite direction. Because the restriction endonucleases *Hind*III and *Eco*RI are between *bar* and *SbSNAC1*, there would be two bands after digestion with these two endonucleases. Meanwhile, for the genome digested with *Dra* I and *Bgl* II with no sites in the insertion sequence, there was only a single band in the Southern blot results. The special structure of the insertion sequence explained the results of Southern blot. Because of the special structure of the insertion sequence, neither the Sanger sequencing method nor the second generation sequencing method could obtain the cloned sequence.

## Discussion

Detailed molecular characteristics of flanking sequences of insertions play an important role in safety assessment of genetically modified crops [30]. Traditionally, the PCR-based methods such as Tail-PCR and genome walking were used to determine locations of integration sites and junction sequences between exogenous sequences and host genome [9, 31]. With the continuous improvement of technology, the flanking sequence of single T-DNA copy insertion transgenic lines can be obtained quickly and cheaply by these PCR-based methods. Charles et al. amplified the 5’ flanking sequence of insertion sequence of 75 *Mu* maize mutant lines based on the PCR method, but the flanking sequences of 20% of the lines could not be obtained by this method in their study [32]. These PCR-based methods may not work well if the deletion, modification or rearrangement occurred in exogenous insertion sequences. On the other hand, high level of duplication or repetitive genome sequences adjacent to the exogenous fragment insertion location might increase the difficulty of identifying the flanking sequences. The maize genome size is about 2.3-2.5 G and nearly 85% of the maize genome is composed of hundreds of families of transposable elements, dispersed unevenly across the whole genome [33, 34]. In our research, the genome walking method was also used to amplify the flanking sequence of the insertion sequences, but only one end of the flanking sequence was identified due to the complex structure of the insertion sequences. As a result, using the PCR-based methods to identify the flanking sequences of complex exogenous fragments of transgenic lines might be a challenge in the maize genome.

With the emergence and development of high throughput next generation sequencing (NGS) technology, the cost of whole genome sequencing has been greatly reduced (Table 4). The NGS technology has been widely used in different species to discover genome structural variation, rearrangement, and so on [35-37], with some advantages including high throughput, no need for large amounts of DNA, time and labor saving [38]. Compared with other methods, the WGS combined with targeted bioinformatics analysis has become a sensitive and efficient method for identifying molecular characteristics of GM crops. Guo et al. used the WGS technology to sequence and analyze the sequence information of two GM soybean events, and successfully identified from one single read analysis [6]. In the work of Kiran et al., by using the NGS method together with the PCR amplification to identify the T-DNA insertion site and flanking sequence of the GM maize IE09S034 at the 3’ end [18]. Although several NGS-based methods have been developed to identify the molecular characteristics of genetically modified crops, some examples often fail to identify insertion sites and flanking sequenced in GM crops. Park et al. used the WGS technology to identify the flanking sequences of three GM rice materials, but one failed to identify the flanking sequences of GM rice [39]. The authors of this article points out that if they can get a longer reads, this problem may not arise. The same problem has arisen in the course of our research. We used the WGS method to sequence the transgenic maize lines. After detailed analysis, only the one end flanking sequence of the insertion fragments was identified. Generally, the NGS technology using to identify the flanking sequences might be efficient if the transgenic line has one or two copies of insertion or stacked transgenic events. On the other hand, the clean reads of the WGS technology are usually only about 150 bp, and assembling the flanking sequences requires a large number of reads in the insertion region to be spliced together, which is a huge challenge for the genome with a large number of repetitive sequences. In our study, ∼64.57 × coverage of the maize genome were sequenced, and only one end flanking sequence was identified. The insertion sequence consisted of two copies of T-DNA sequences, and it had no further clear sequence information, which increased the difficulty of identifying the flanking sequence using the WGS technology. Increasing the sequence coverage by deep sequencing might be helpful to identify the other end flanking sequence. But it is still difficulty to characterize the structure of the exogenous sequences using the WGS technology.

**Table 4.** Characteristic comparison of three methods for obtaining flanking sequences.

| Method | Time | Cost | Insertion sequence structure | Genome complexity | Flux level |
| --- | --- | --- | --- | --- | --- |
| PCR-based | Short | Cheap | Simple | Simple | Low |
| NGS | Short | Cheap | Simple | Simple | High |
| Fosmid+sequencing | Long | Expensive | Complex | Complex | Low |

The fosmid technology has been applied in genomics of many species, such as rice [40], maize [41], and human [42]. Compared with the BAC library, the construction of fosmid library is simpler and faster. Furthermore, average length of the insertion sequences of the fosmid library is 38-48 kb which might be suitable to identify the flanking sequences and characterize the structure of the insertion sequence of transgenic lines. On the other hand, read length of single-molecule real-time DNA sequencing might be 10-20 kb, which may also contribute to characterize the insertion sequence of transgenic events. In our study, three positive clones were accurately identified from the fosmid library using the PCR method with three pairs of specific primers. Furthermore, with the SMRT sequencing technology, the flanking sequences were identified and the structure of exogenous insertion sequences was characterized. Although the use of the method of building fosmid libraries and the third generation sequencing to obtain flanking sequences of GM crops is more time-consuming and costly than the method based on PCR and WGS, it is more reliable for some GM crops with complex genomic or insertion sequence structure. In identifying the flanking sequences of GM crops, the method of constructing fosmid libraries combined with the third-generation sequencing technology is not a high-throughput method, it is more time-consuming and costly, but it is more reliable to effectively identify the flanking sequences and characterize the insertion sequences with deletion, modification or rearrangement. As a result, when identifying the flanking sequences of genetically modified crops, different methods should be flexibly selected according to their genomic characteristics and the internal structure of insertion sequences (Table 4).

## Supporting Information

**S1 Figure. Vector for transgenic line.**

**S2 Figure. Electrophoretogram of fosmid clones digested with *Not I***. 1-16: Insert fragments; M: marker.

## Acknowledgments

This work was carried out with the support of Innovation Program of Chinese Academy of Agricultural Sciences and the Major Projects of Genetically Modified Organisms, China (2016ZX08003004).

